# Real-Time Interactive Visualization and Analysis of Neurotransmitter Data

**DOI:** 10.1101/2022.09.03.506483

**Authors:** Anne Thomas Homescu, Teresa Murray

## Abstract

We describe an interactive visualizer (implemented in R Shiny framework) to facilitate analysis and a better understanding of neurotransmitter data collected within the context of epileptic seizures.

Given the very high granularity of collected data (at millisecond level), it is challenging to use static visuals and/or tables for deeper data insights and features. Such challenges are greatly alleviated through an interactive visualizer (dashboard) which has ability to zoom out (for “big picture” analysis) and to zoom in (for a much more focused and targeted targeted analysis).

The visualizer is available at link https://kittyviz.shinyapps.io/GluGabaViz

## 1 Introduction

Epilepsy is one of the most prevalent neurological disorder affecting more than 50 million people worldwide Pitkanen et al. (2016), Koshal et al. (2018) and an estimated 2-4 million people are newly diagnosed with epilepsy each year. However, despite over 20 anti-epileptic drugs being available, seizures remain uncontrolled in about 30% of patients Pitkanen et al. (2016).

One of the major factors that have impeded rapid progress is the complex and multifactorial nature of epilepsy, and its heterogeneity. Therefore, the vision of developing targeted treatments for epilepsy is more and more relying upon the development of biomarkers that allow individually tailored treatment Pitkanen et al. (2016).

A number of molecular biomarkers for epilepsy have been identified including glutamate (GLU), *γ*-aminobutyric acid (GABA), and miRNAs Thompson and Ackroyd (2020). Glutamate and GABA are the major and most abundant neurotransmitters in the brain and contribute to signal transmission as excitatory or inhibitory drives, respectively Çavdar et al. (2019). Numerous studies have indicated that an imbalance in glutamatergic (excitatory) and GABAergic (inhibitory) neurotransmitter system is one of the dominating pathophysiological mechanisms underlying the occurrence and progression of seizures. Further, this alteration in GABAergic and glutamatergic system disrupts the delicate balance of other neurotransmitters system in the brain Koshal et al. (2018). Past results from patients demonstrated that glutamate levels were higher in epileptic tissue both interictally and during seizures. In contrast, GABA was shown either to increase or decrease before seizures. Further research is necessary to determine how these molecules affect the pathogenesis of epilepsy Luna-Munguia et al. (2019).

Batra et al. (2014) published research on the fabrication of a device for the detection of GLU. The focus of this work was to detect glutamate in food, while pointing out the importance of the molecule in epilepsy Thompson and Ackroyd (2020). However, no real-time seizure data was presented. Cordeiro et al. (2015) developed and characterized an implantable multiplex microbiosensor for in vivo in real time simultaneous monitoring of the three major biomarkers in brain energy metabolism: glucose lactate and pyruvate. While it was mentioned that t this system has potential use for epileptic diagnosis Thompson and Ackroyd (2020), no real-time seizure data was presented. Ganesana et al. (2019) reported the development of a novel glutamate biosensor for in vivo monitoring. While the study was mentioning its epilepsy applicability, no specific seizure data was presented Thompson and Ackroyd (2020).

GABA is a major inhibitor neurotransmitter that helps with plasticity and network synchronization in the brain. To determine the continuous concentration of GABA, Hossain et al. (2018) developed a probe capable of detection at two sites. On the first a Pt electrode, only GOx is present, while the second Pt electrode has both GOx and GABASE. Both were coated with glutaraldehyde and BSA, as well as m-phenylenediamine. The amount of GABA present was determined from the difference of the H2O2 current at each sensor: IGABA = IH2O2(Site 2) - IH2O2(Site 1). What is unique about this biosensing device is that no extra reagents were required to achieve a signal, since in situ *α*-ketoglutarate is a product of glutamate oxidation. The key reaction will be continuous in vivo because glutamate is readily available. This biosensor was able to measure GABA with 26 times more sensitivity than any other reported in literature so far, at 36 ± 2.5 pA *μ*M–1 cm–2 with an LOD of 2 *μ*M in vitro, while the GLU results are only comparable. In ex vivo experiments of the rat hippocampal brain slices, during stimulations of the rat brain, GLU concentrations ranged from 5 to 25 *μ*m, and GABA ranged from 5 to 13 *μ*M Thompson and Ackroyd (2020).

These results demonstrate that the developed GABA probe constitutes a novel and powerful neuroscientific tool that could be employed in the future for in vivo longitudinal studies of the combined role of GABA and Glu (a major excitatory neurotransmitter) signaling in brain disorders, such as epilepsy and traumatic brain injury, as well as in preclinical trials of potential therapeutic agents for the treatment of these disorders Hossain et al. (2018).

As noted above, dysregulation of glutamate signaling is associated with severe neuropathological conditions, such as epilepsy. Glutamate signals are currently detected by several types of neurochemical probes ranging from microdialysis-based to enzyme-based carbon fiber microsensors. However, an important technology gap exists in the ability to measure glutamate dynamics continuously, and in real time, and from multiple locations in the brain, which limits the ability to further understand the involved spatiotemporal mechanisms of underlying neuropathologies. To overcome this limitation, Scoggin et al. (2019) developed an enzymatic glutamate microbiosensor, in the form of a ceramic-substrate enabled platinum microelectrode array, that continuously, in real time, measures changes in glutamate concentration from multiple recording sites. The probe was able to measure 10-570 *μ*M Glu in vitro, with a sensitivity of 62.3 ± 6.1 nA/*μ*M cm2 in basal media and 270 ± 28 nA/*μ*M cm2 in PBS buffer, which is nearly four times better than other Pt-MEA Glu biosensors Thompson and Ackroyd (2020).

These results confirm that the developed glutamate microbiosensor array can become a useful tool in monitoring and understanding glutamate signaling and its regulation in normal and pathological conditions Scoggin et al. (2019).

Most studies so far have only looked at GLU and GABA in isolation as biomarkers for epilepsy. However, a more meaningful interpretation of the mechanisms of disease and secondary injury can be obtained by knowing the regional E/I balance. Doughty et al. (2020) presented a novel microwire-based biosensor probe for simultaneous real-time measurement of glutamate and GABA dynamics in vitro and in vivo. Toward their long-term goal of longitudinal, in vivo recording, the researchers recorded GABA and GLU from a freely moving rat at 2, 10, and 16 weeks after cannula implantation using different microwire biosensors for each session. Changes in E/I and the pattern of fluctuations in neurotransmitter levels during sleep, awake behavior, and epileptic signaling were consistent with reports in the literature Doughty et al. (2020), thus further substantiating E/I ratio as a potential useful biomarker for epilepsy.

## 2 Data description

Data used in this visualizer was obtained using procedures described in Doughty et al. (2020). See details for dataset used in Fig 8 (of that study) which presented how in-vivo GLU and GABA dynamics change with different types of behavior. Current shown in all panels is after subtraction of sentinel current for both channels and subtraction of GLU current from the GABA channel.

**Figure 1:**
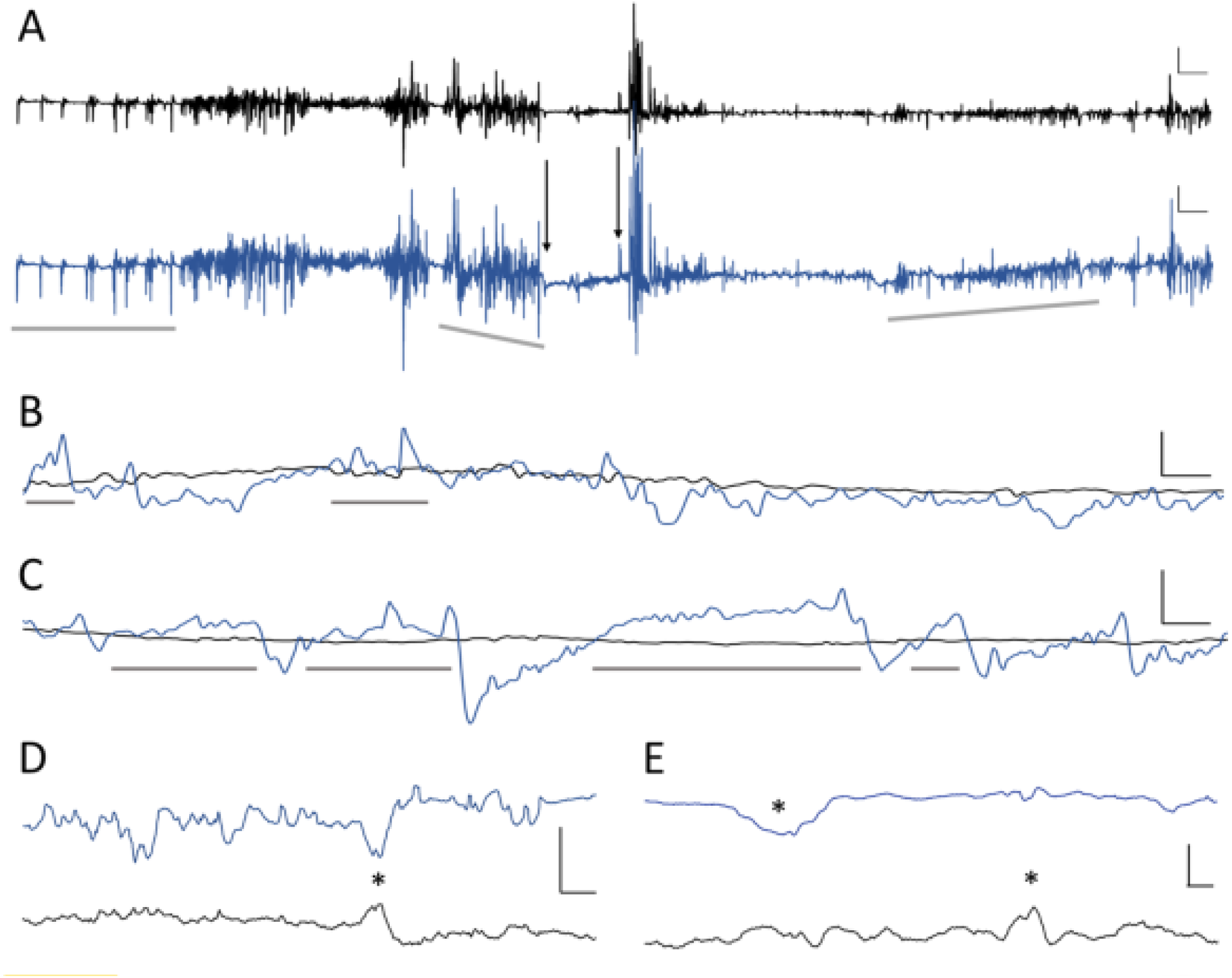
In vivo dynamics: GLU (black) and GABA (blue) Source: Doughty et al. (2020)

**Figure 2:**
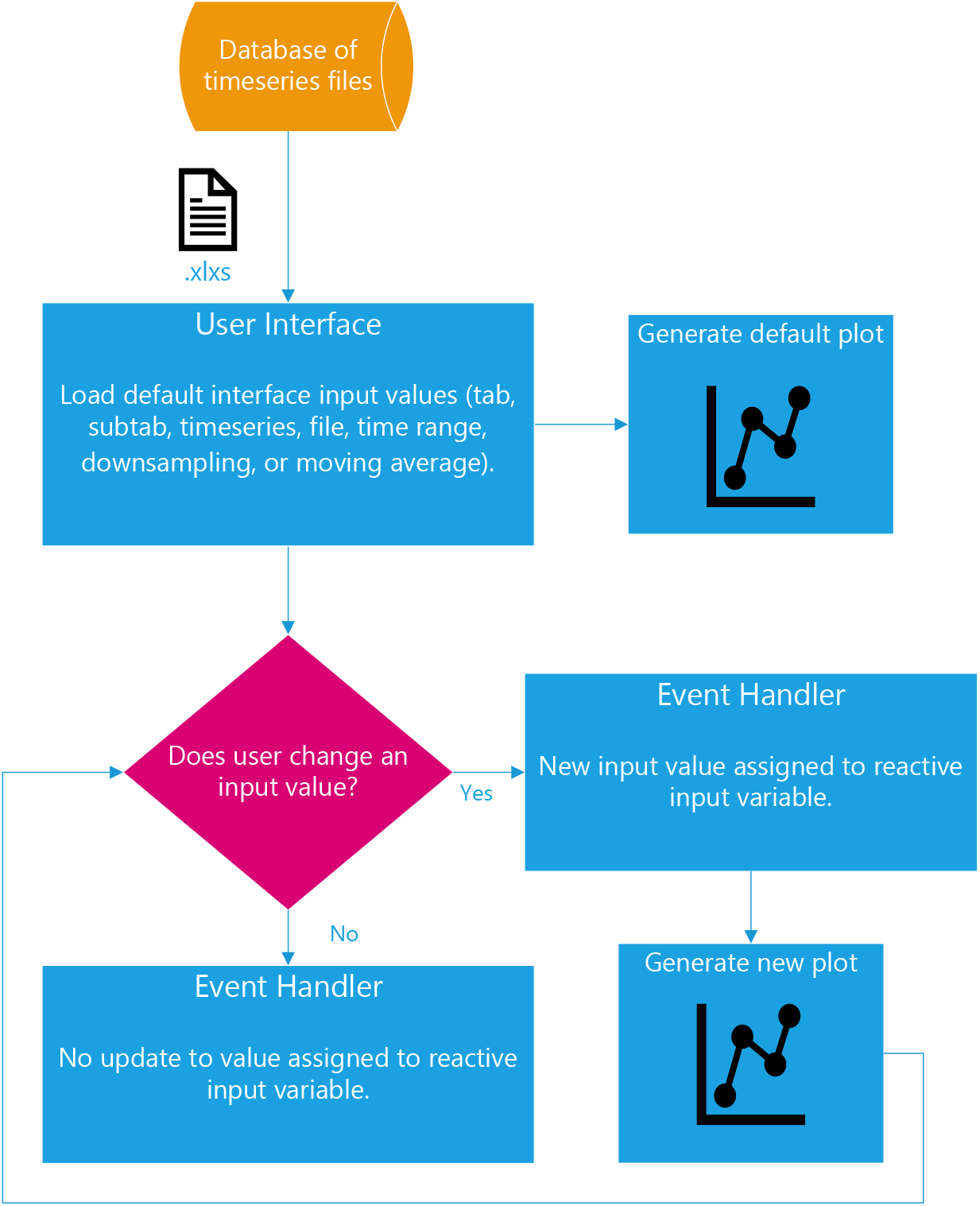
Flowchart of Shiny dashboard

**Figure 3:**
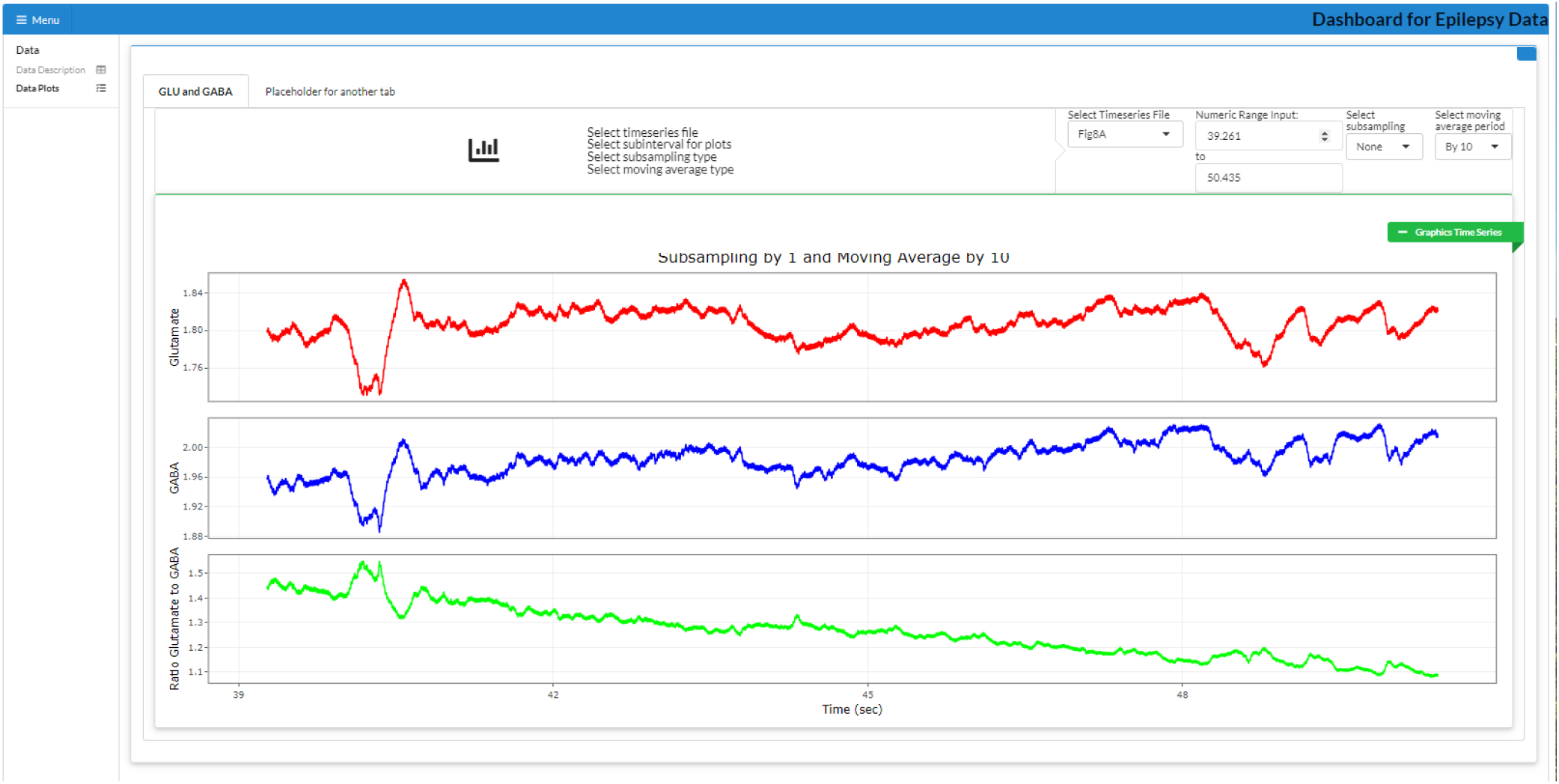
Dashboard interface - Glutamate (top); GABA (middle); Ratio (bottom).

**Figure 4:**
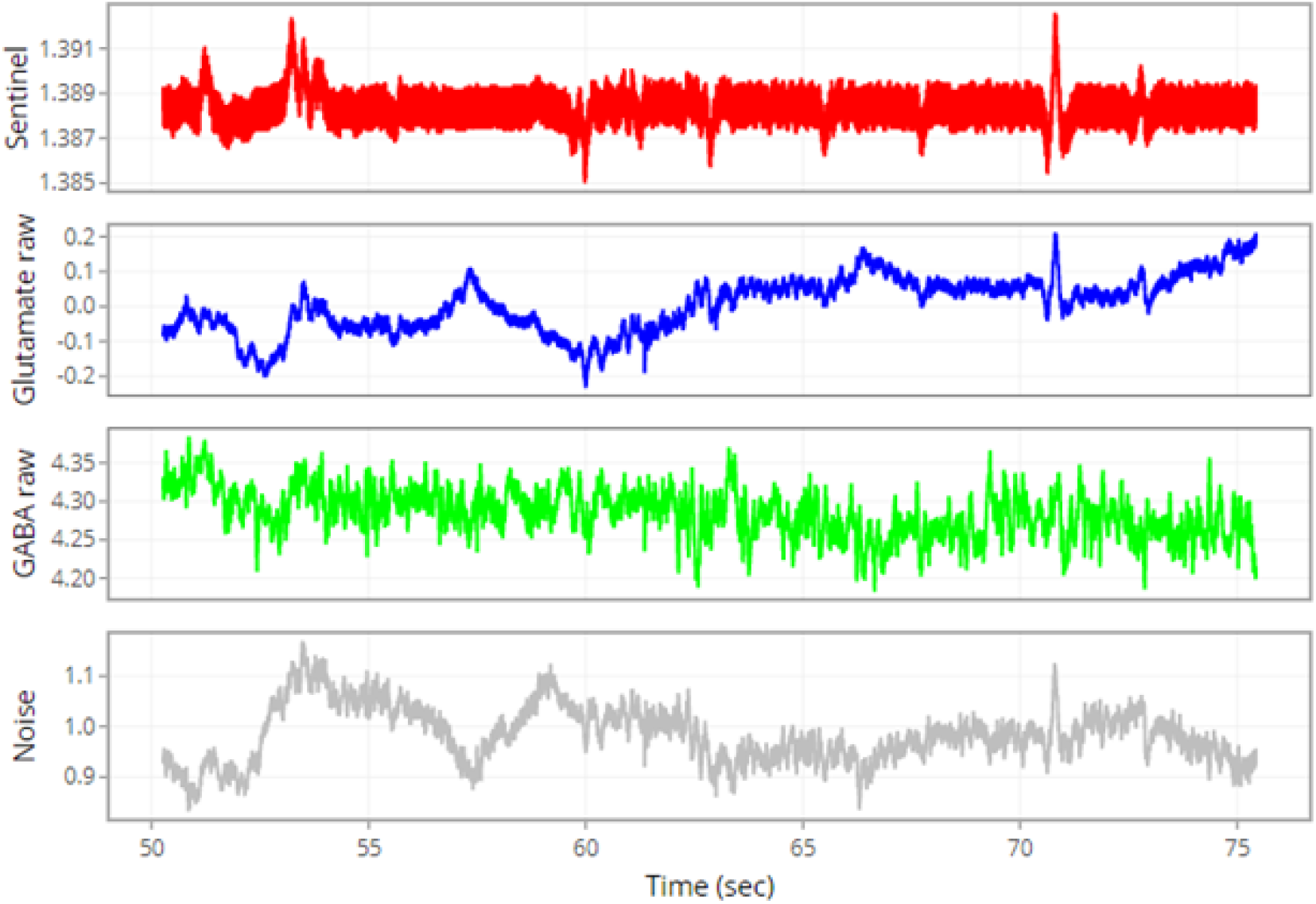
Plot of raw data for all channels.

**Figure 5:**
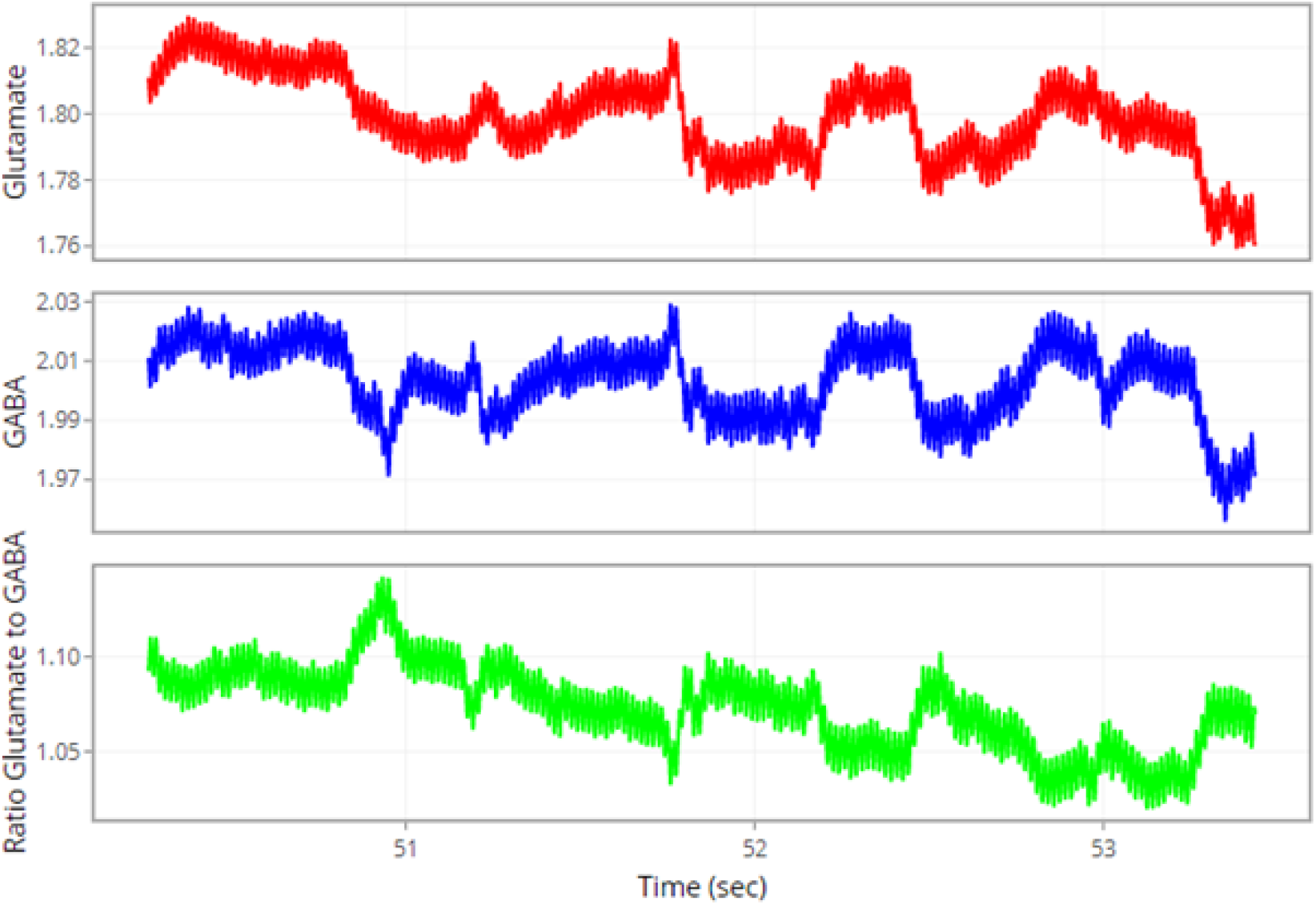
Plots of adjusted data for all channels

**Figure 6:**
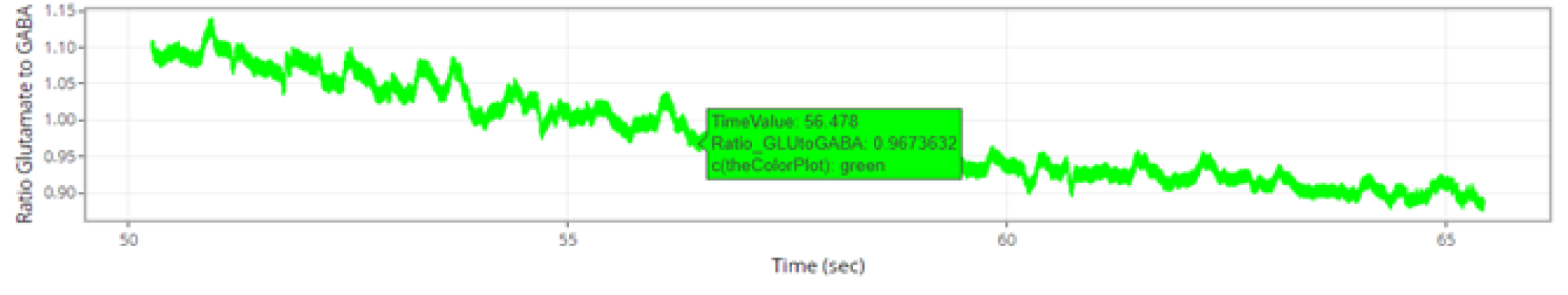
Cursor and value of a datum

**Figure 7:**
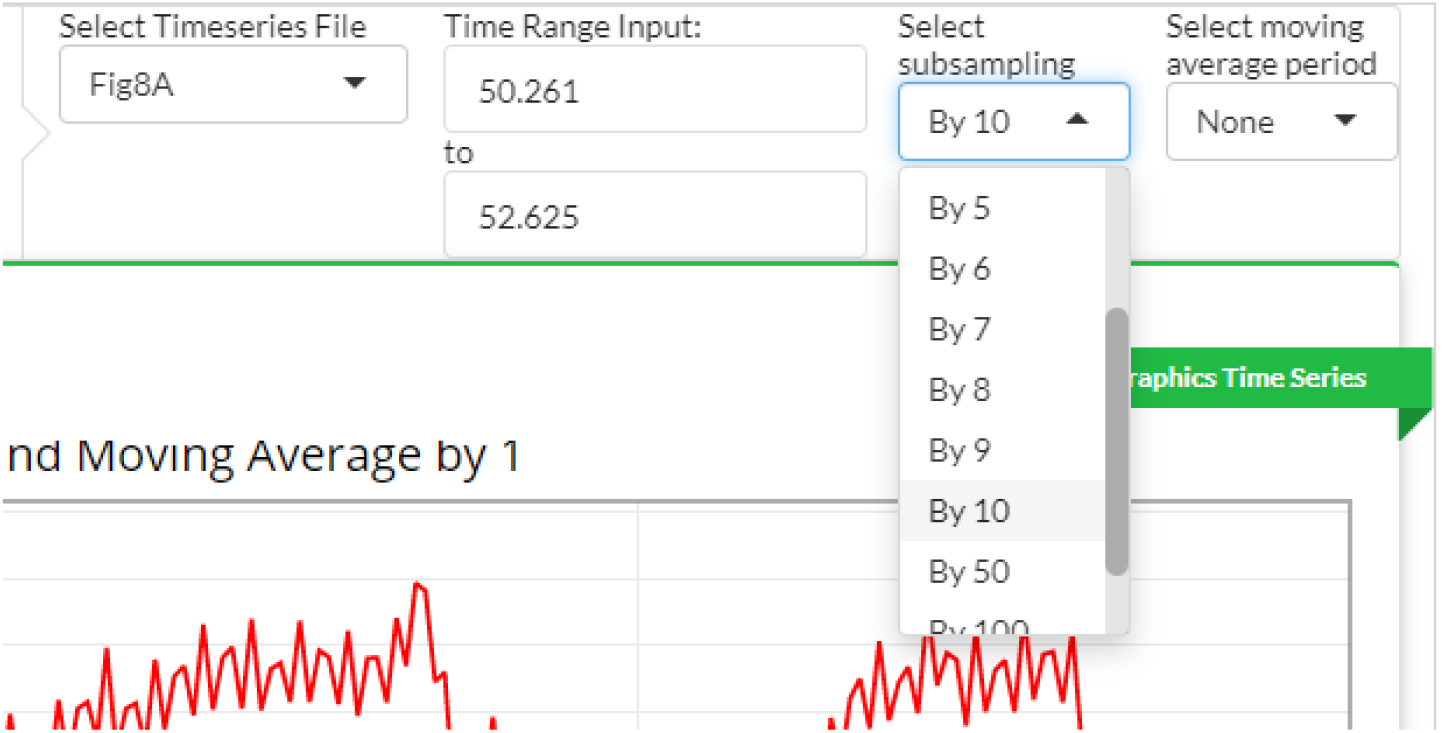
Example of drop-down menu

**Figure 8:**
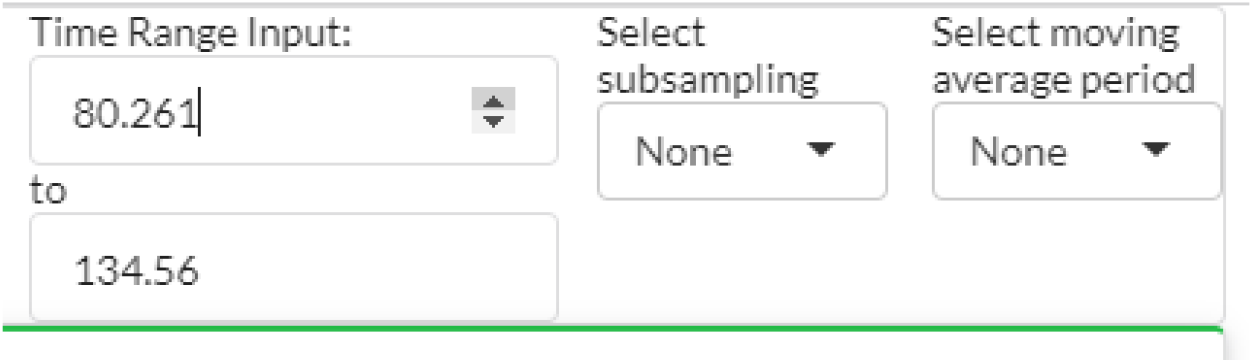
Example of scroll menu

Several behaviors are captured in the 8-min 40-s recording shown in Panel A. Interictal-like peaks appear in the first 70 s when the rat is not moving. This is followed by walking and grooming. Movement ceased between the two downward arrows as the rat abruptly stopped moving. A rapid decrease in GLU and GABA occurred at the onset of freezing. GABA gradually increased until the point when the rat clearly exhibited seizure behavior with forelimb clonus, which is a Racine Scale 3 behavior during epileptic seizures in rats. After this, GABA and GLU fluctuated rapidly, and after a few seconds the rat reared and fell, which are Racine Scale 4 and 5 behaviors, respectively. Maximum current fluctuations in both biosensor channels were observed just before and through rearing. After the rat ceased rearing, it continued to exhibit mouth and facial movements with some occasional head bobbing for the remainder of this recording segment.

Panels B and C visualized representative sleep-wake cycles at Wk 16 that illustrate the changes in GABA. GABA current steadily increased prior to each of six episodes of sleep, during the rat’s light cycle, and remained elevated with some fluctuation during sleep. Immediately prior to waking, GABA levels rapidly fell and remained low during activity.

Panels D and E visualized representative recording two (respectively eight) weeks after cannula implantation while the rat was walking in its home cage.

Vertical and horizontal scale bars represent different time scales:

- Panel A: vertical scale bars correspond to 1 nA while horizontal scale represents 20 s for both GLU and GABA traces
- Panel B: vertical scale bars correspond to 0.5 nA while horizontal scale represents 2 min
- Panel C: vertical scale bars correspond to 1.0 nA while horizontal scale represents 2 min
- Panel D: vertical scale bars correspond to 0.1 nA while horizontal scale represents 125 ms
- Panel E: vertical scale bars correspond to 0.2 nA while horizontal scale represents 100 ms

The tool described in the next section is intended to aid in further analysis of the in vivo recordings presented in Doughty et al. (2020). This dataset is considered throughout Section (4.3) and Section (4.3) of this article. For consistency with Doughty et al. (2020), this database is denoted as “Fig. 8” in the dropdown menu of the interactive visualizer described in this report.

## 3 The need for an interactive visualizer

While the benefits and importance of static and interactive visualizations have been explored heavily by researchers, it is also important to note the difference between the two. Static visualizations graphically present findings that do not change over time, hence only depict the situation or viewpoint at the time they were generated. On the contrary, interactive visualizations allow the user to explore, interact and directly control the graphical representations generated; this is usually done by allowing the user to modify parameters that are mapped to underlying configurations applied on the data or models used for the visualizations. Interactive visualizations are beneficial in various ways such as allowing for more than one viewpoint to be generated, unlike the single viewpoint in static visualizations, and effective and efficient identification of relationships and trends as the user can change parameters and instantly observe their impacts. In addition, interactive visualizations can simplify and present complex data, and assist in dynamic data storytelling, which is facilitated by changing parameters and exploring multiple viewpoints. Nonetheless, static visualizations are also important and useful when dealing with less complex data stories, or when the requirement is to focus on portraying a predetermined view rather than exploration. Understanding the difference between the two approaches is important, as the selection of either approach should be based on the purpose of the visualization to avoid over- or under-engineering the solution and failing to achieve the expected outcomes Khedr and Hilal (2021).

R (together with Shiny) and Python (together wih streamlit, plotly, bokeh, etc.) are widely recognized to the forefront as open-source, powerful tools for creating interactive visualizations, and data visualization can be efficiently done through R and Python packages. Both R and Python are open source and cross-platform, and offer a large standard library of well-documented functions, operators and toolkits, as well as over 15,000 packages (for both R and Python) to implement graphical user interfaces (GUIs), automation, text/image processing, and scientific computing Curtis et al. (2019)

While we have selected R/Shiny for our dashboard, we would also like to mention software products for real-time analysis of time series sensor data designed with Python in mind.

**Brainstorm** is a collaborative open-source application dedicated to magnetoencephalography (MEG) and electroencephalography (EEG) data visualization and processing, with an emphasis on cortical source estimation techniques and their integration with anatomical magnetic resonance imaging (MRI) data. The primary objective of the software is to connect MEG/EEG neuroscience investigators with both the best-established and cutting-edge methods through a simple and intuitive graphical user interface (GUI) Tadel et al. (2011). Brainstorm provides a rich interface for displaying and interacting with MEG/EEG recordings (Figure 2) including various displays of time series (a)–(c), topographical mapping on 2D or 3D surfaces (d)-(e), generation of animations and series of snapshots of identical viewpoints at sequential time points (f), the selection of channels and time segments, and the manipulation of clusters of sensors Tadel et al. (2011). Although Brainstorm was built from a vast amount of pre-existing lines of Matlab code as its methodological foundations for data analysis, the authors acknowledge that Python might be a better choice for a new project because of its non-commercial open source license Tadel et al. (2011).

Software for the Analysis and Continuous Monitoring of Electrochemical Systems (**SACMES**) is an open-source, easily customizable, multiplatform compatible program for the real-time control, processing, and visualization of large volumes of electrochemical data. The software’s architecture is modular and fully documented, allowing the easy customization of the code to support the processing of voltammetric (e.g., square-wave and cyclic) and chronoamperometric data. The program also includes a graphical interface allowing the user to easily change analysis parameters (e.g., signal/noise processing, baseline correction) in real-time Curtis et al. (2019).

## 4 Interactive Visualizer (Dashboard)

The importance of analytics and visualization tools has been growing over the last decades to handle big data which stems from all aspects of life. Data visualization is the presentation of data in a pictorial or graphical format, and a data visualization tool is the software that generates this presentation. Data visualization provides users with intuitive means to interactively explore and analyze data, enabling them to effectively identify interesting patterns, infer correlations and causalities, and supports sense-making activities Bikakis (2018). The importance of analytics and visualization tools has been growing over the last decades to handle big data Khedr and Hilal (2021). Visualization is the best way to analyze data, and interactive visualization is even better as it allows the user to interact with data in real-time.

### 4.1 Shiny framework for interactive visualization

Our application is implemented in R/Shiny, the details of which are described in the next sub-section. As such, the remainder of this section focuses on the Shiny package for R, have allowed R programmers to interactively show the output for R programs to Web-browsers.

Though there are several tools that allow interactive visualization such as Tableau or Microsoft’s Power BI, the combination of Shiny and R has been selected for R’s powerful statistical backend which includes ready-made packages with various statistical algorithms and the ability to leverage Shiny’s extensive and customizable frontend capabilities to present a customized interactive web applications tailored for the specific requirements Khedr and Hilal (2021). Shiny allows customization of the application’s user-interface to provide an elegant environment for displaying user-input controls and simulation output–where the latter simultaneously updates with changing input Wojciechowski et al. (2015).

The R Shiny platform offers six distinct advantages: (1) the user interfaces that one can develop through R Shiny are easy to use and do not require any a priori training, (2) R Shiny is freely accessible and does not require a license, and (3) because R Shiny is based in the R statistical programming language, statistical modeling capabilities can be seamlessly integrated within the web application, including novel and recently published statistical methods with corresponding R packages that contain functions to implement such methods Xia et al. (2022), (4) Shiny Web-browser interface that can be viewed on the localhost (user’s own computer) or on another computer accessed by means of the Internet, (5) Applications have been developed using the Shiny package and R and can be viewed without an R installation or files containing R code, (6) Unlike other Web-page design methods, only previous experience with the R programming language is required Wojciechowski et al. (2015).

Development of interactive web applications to deposit, visualize and analyze biological datasets is a major subject of bioinformatics. R is a programming language for data science, which is also one of the most popular languages used in biological data analysis and bioinformatics. However, building interactive web applications was a great challenge for R users before the Shiny package was developed by the RStudio company starting from 2012. By incorporating HTML, CSS and JavaScript code into R code, Shiny has made it incredibly easy to build web applications for the large R community in bioinformatics and for even non-programmers [Example of Shiny](https://academic.oup.com/bib/article-abstract/23/1/bbab415/6387320).

For example, Salehi et al. (2021) provides a useful online interactive [dashboard](https://mahdisalehi.shinyapps.io/Covid19Dashboard) to visualize and follow confirmed cases of COVID-19 in real-time. This dashboard is intended as a user-friendly dashboard for researchers as well as the general public to track the COVID-19 pandemic, and is built in open-source R software (Shiny in particular). The R Shiny framework serves as a platform for visualization and analysis of the data, as well as an advance to capitalize on existing data curation to support and enable open science Salehi et al. (2021).

MEPHAS is a free interactive GUI of R that was developed to support statistical data analyses for medical and pharmaceutical students and practitioners. Through a shiny framework, MEPHAS provides a web-based interactive GUI that can dynamically visualize real-time analytical results across multiple panels simultaneously. Notably, this analytical mode avoids repetitive and redundant operations when finding optimal parameters, and therefore, the analytical process is facilitated. This interactivity enables users to view the real-time output and adjust all parameters during the analytical process, saving much time that was spent in repetitive operations with static GUI panels in other software Zhou et al. (2020).

Xia et al. (2022) introduced a user-friendly, interactive web application, shinyOPTIK, to improve the usability of the OPTIK database and in doing so, facilitate a better understanding of cancer risk factors and mortality trends across the KU Cancer Center catchment area.

TPWshiny is a standalone, easy to install, R application to facilitate more interactive data exploration without requiring programming skills. TPWshiny provides an intuitive and comprehensive graphical interface to help researchers understand the response of tumor cell lines to therapeutic agents. The data are presented in interactive scatter plots, heatmaps, time series and Venn diagrams. Data can be queried by drug concentration, time point, gene and tissue type Zhang et al. (2021).

EDTox is a user-friendly Shiny application that allows non-expert R users to utilize large-scale toxicogenomics data for the identification and prioritization of endocrine disruption compounds Sakhteman et al. (2022).

We leveraged the Shiny framework in combination with R packages to develop our interactive dashboard for real-time display and analysis of large-volume neurochemical data. The interface allows the user to dynamically adapt analysis parameters for real-time contraol and visualization.

### 4.2 Implementation

We describe the user-friendly and interactive web application that we have developed to facilitate a better understanding of epileptic seizures. The web-based visualization application was implemented under the R Shiny framework within the RStudio integrated development environment for the R statistical programming language. We have used R version 4.2.1 with packages as of August 1, 2022.

The visualization R packages (such as ggplot2, ggvis, and plotly) enabled us to quickly analyze and visualize data as histograms, bar graphs, and scatterplots, and to customize plots with various themes and coloring options. The user may also create, design, and build interactive dashboards using Shiny.

The Shiny app is deployed on the platform “shinyapps.io”, which hosts each app on its own virtualized server (called an instance) Salehi et al. (2021).

The reader can access the link for Visualizer for Glatmate and GABA data captured in real-time.

### 4.3 Features and capabilities

The user interface UI includes the following features and capabilities:

- ability to select among multiple datasets
- cursor hover over single points
- dropdown menus
- scroll menu for time ranges

#### 4.3.1 Ability to select among multiple datasets

User may switch between tabs and panels to display the specific dataset under analysis.

The ***Channels*** tab displays the separate channel signals:

1. Sentinel
2. Glutamate
3. GABA
4. Noise.

The following figures show

- Fig. 4: raw data for all channels:
- Fig. 5: adjusted data for all channels

♢ GLU channel after sentinel is subtracted
♢ GABA channel after GLU and sentinel are subtracted
♢ ratio GLU/GABA

The ***GLU and GABA*** tab displays the GLU, GABA, and GLU-to-GABA ratio signals with the Sentinel signal subtracted from the GLU and GABA signals.

#### 4.3.2 Cursor hover over single points

User can hover the cursor over a plot datum to to see exact time and value.

#### 4.3.3 Drop-down menus

Each panel includes drop-down menus for selecting a particular dataset (Select Timeseries File), selecting a subsampling value (Select subsampling), and selecting a moving average window (Select moving average period).

#### 4.3.4 Scroll menu for time range

Each panel includes a scroll menu for selecting the time range.

## 5 Analysis

The sub-sampling and moving average menu options allow the investigator to zoom in on specific features at different levels of granularity. We exemplify these capabilities through a second dataset included in the visual dashboard.

Details of this dataset are given in Doughty et al. (2020), which use enzyme-based microelectrode array biosensors present the potential for improved biocompatibility, localized sample volumes, and much faster sampling rates over existing measurement methods. The dataset contains measurements of Glutamate (GLU) and *γ*-aminobutyric acid (GABA) as the major excitatory (E) and inhibitory (I) neurotransmitters in the brain, given that dysregulation of the E/I ratio is associated with numerous neurological disorders.

By measuring real-time dynamics of GLU and GABA in hippocampal slices it was observed that a significant, nonlinear shift in the E/I ratio goes from from excitatory to inhibitory dominance as electrical stimulation frequency increased from 10 to 140 Hz, suggesting that GABA release is a component of a homeostatic mechanism in the hippocampus to prevent excitotoxic damage. This analysis can be enhanced through visualizer capabilities to select different values of subsampling (i.e., downsampling) and of moving average windows in order to obtain information at different granularities.

### 5.1 Subsampling

In the figures below the range of input time is 100.2 to 134.56 seconds, while subsampling is considered at various valuues (10, 50, 100 and 500)

**Figure 9:**
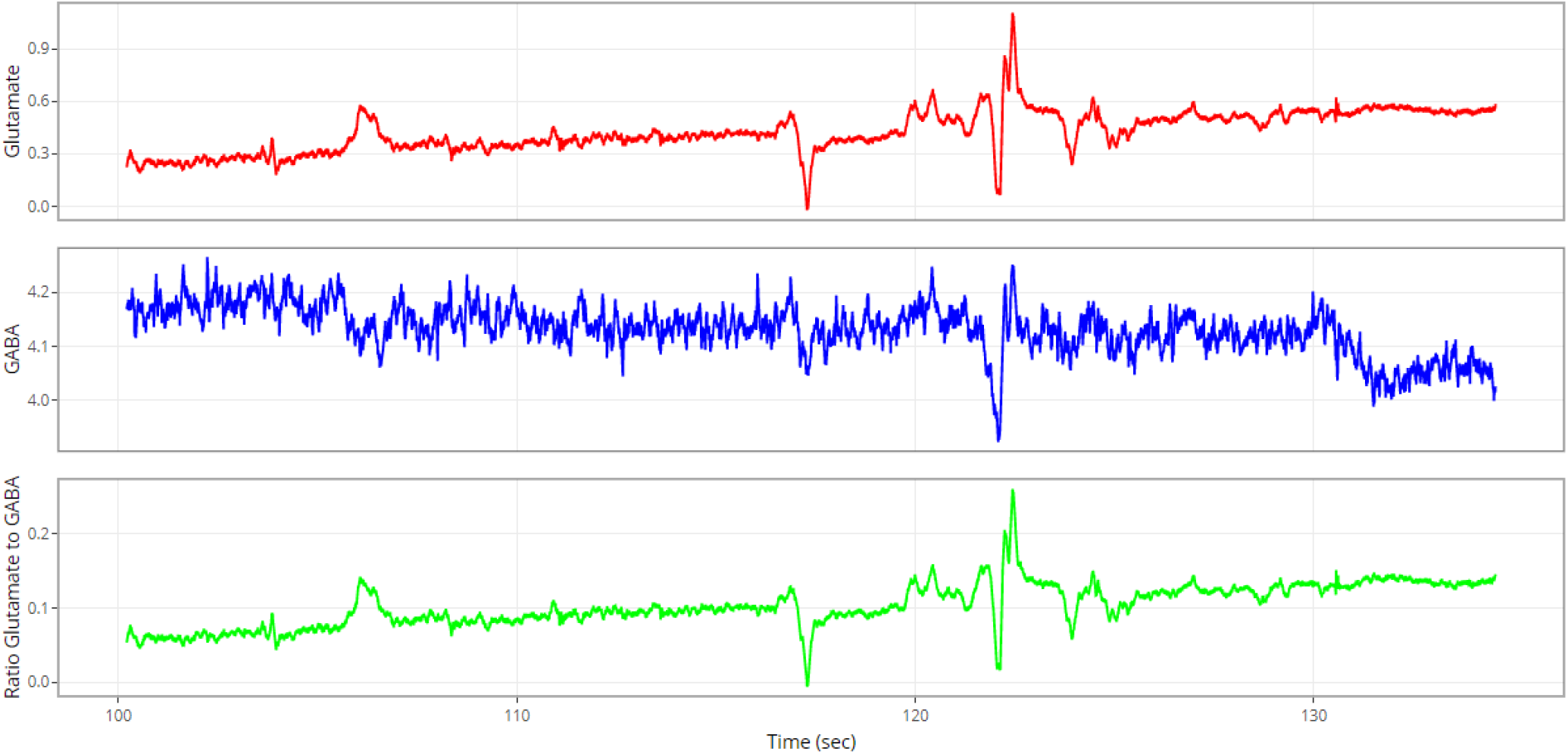
Subsampling by 10.

**Figure 10:**
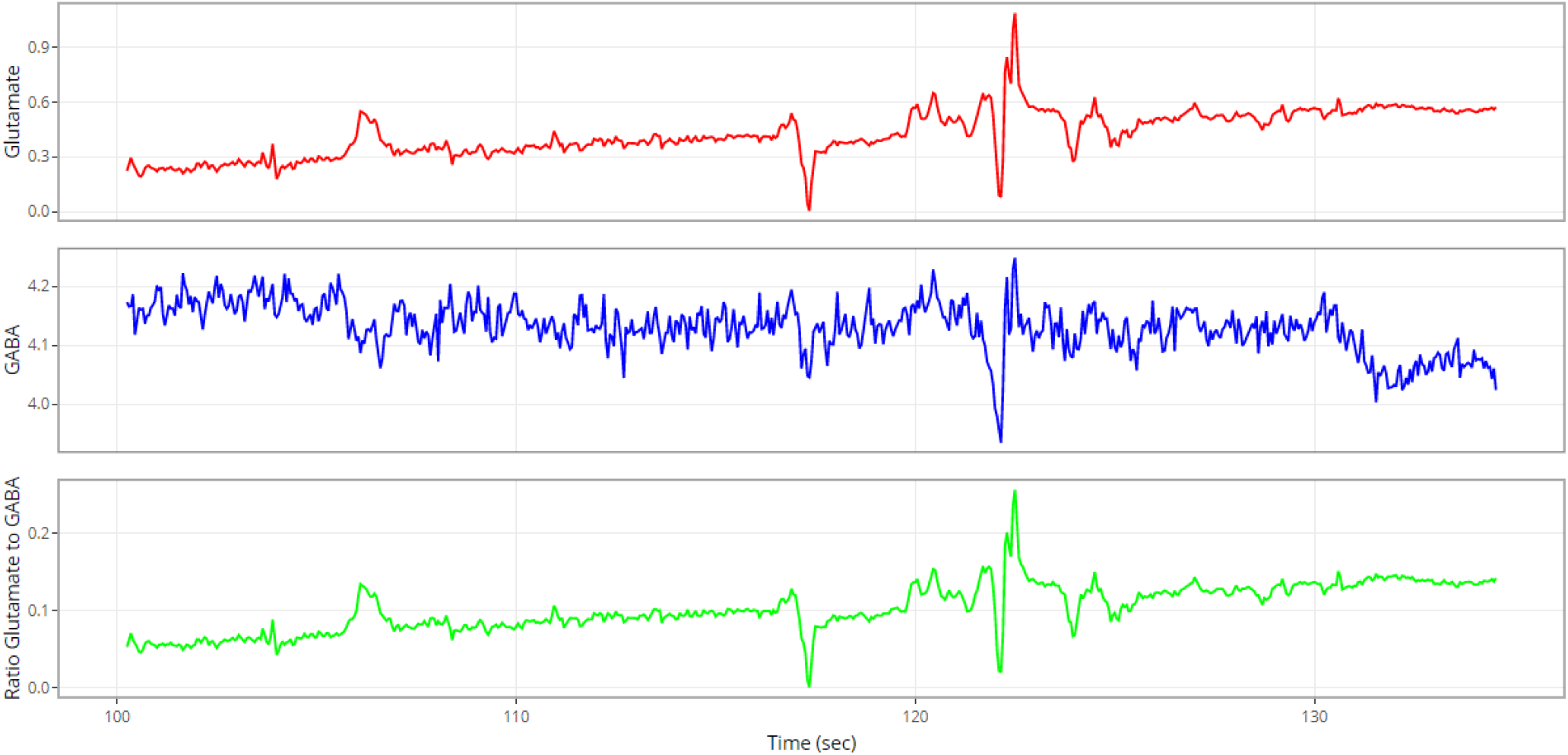
Subsampling by 50.

**Figure 11:**
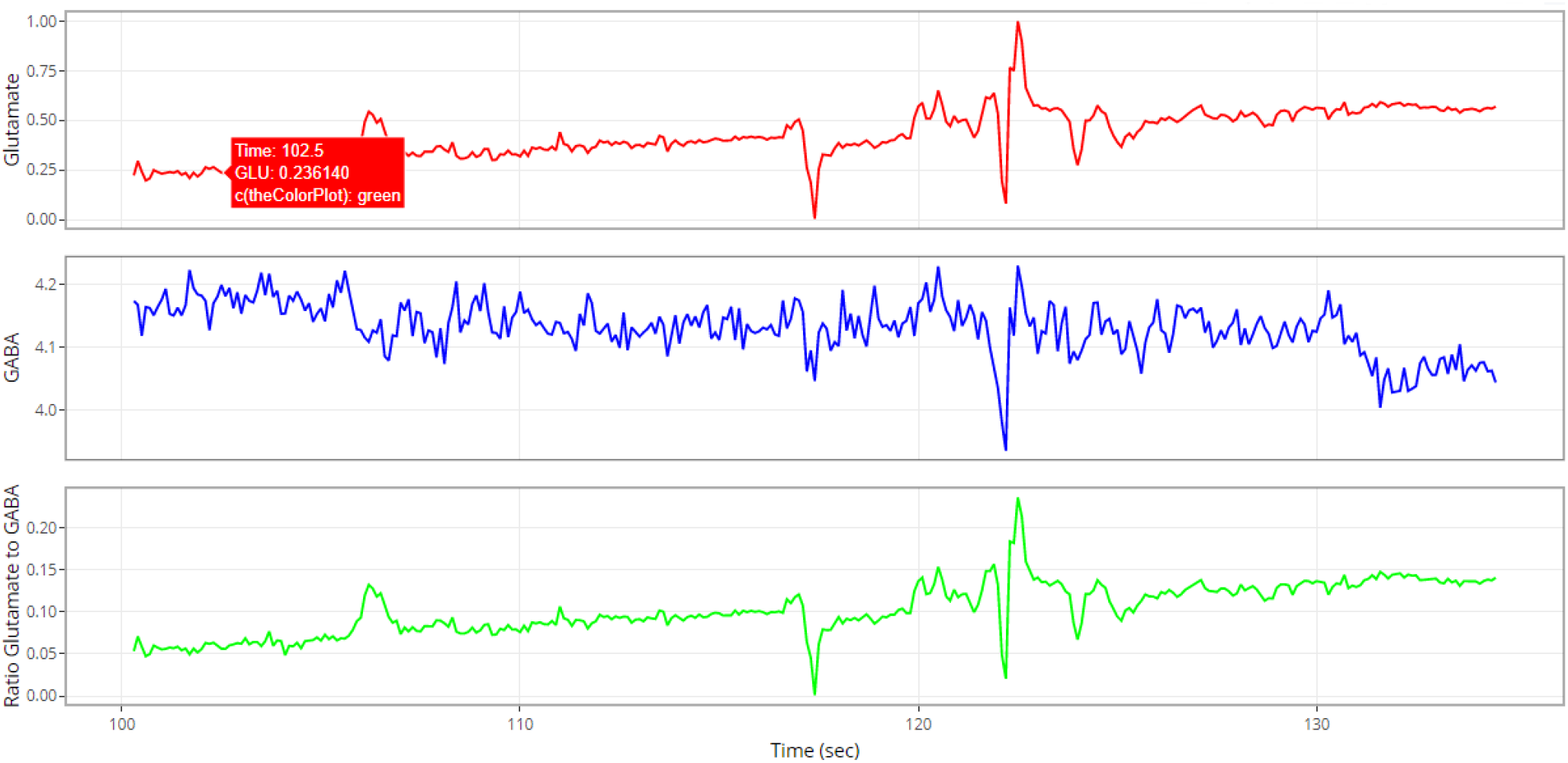
Subsampling by 100.

**Figure 12:**
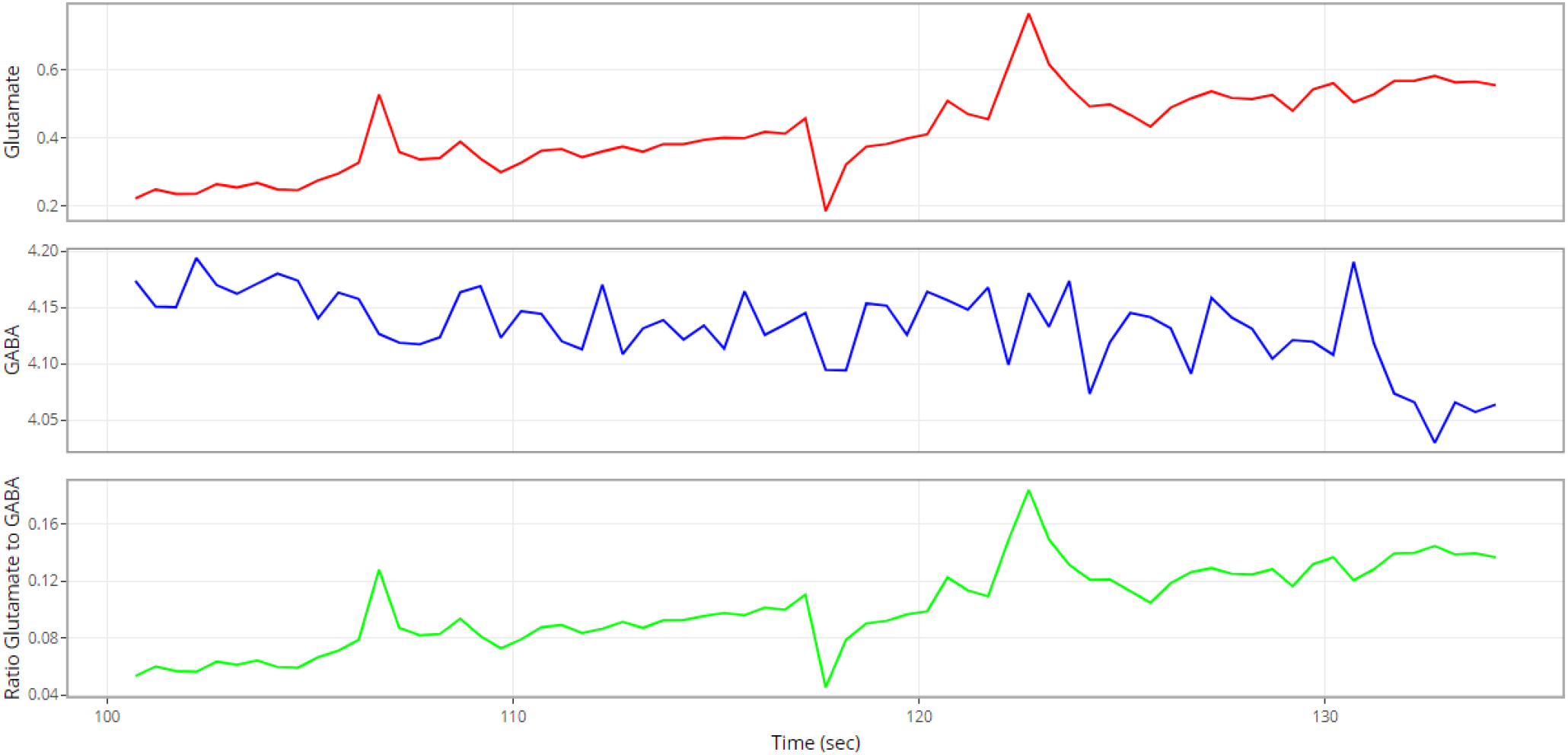
Subsampling by 500.

### 5.2 Moving average window

In the figures below the range of input time is 100.2 to 134.56 seconds, while moving average window is considered at various values (10, 50, 100 and 500).

**Figure 13:**
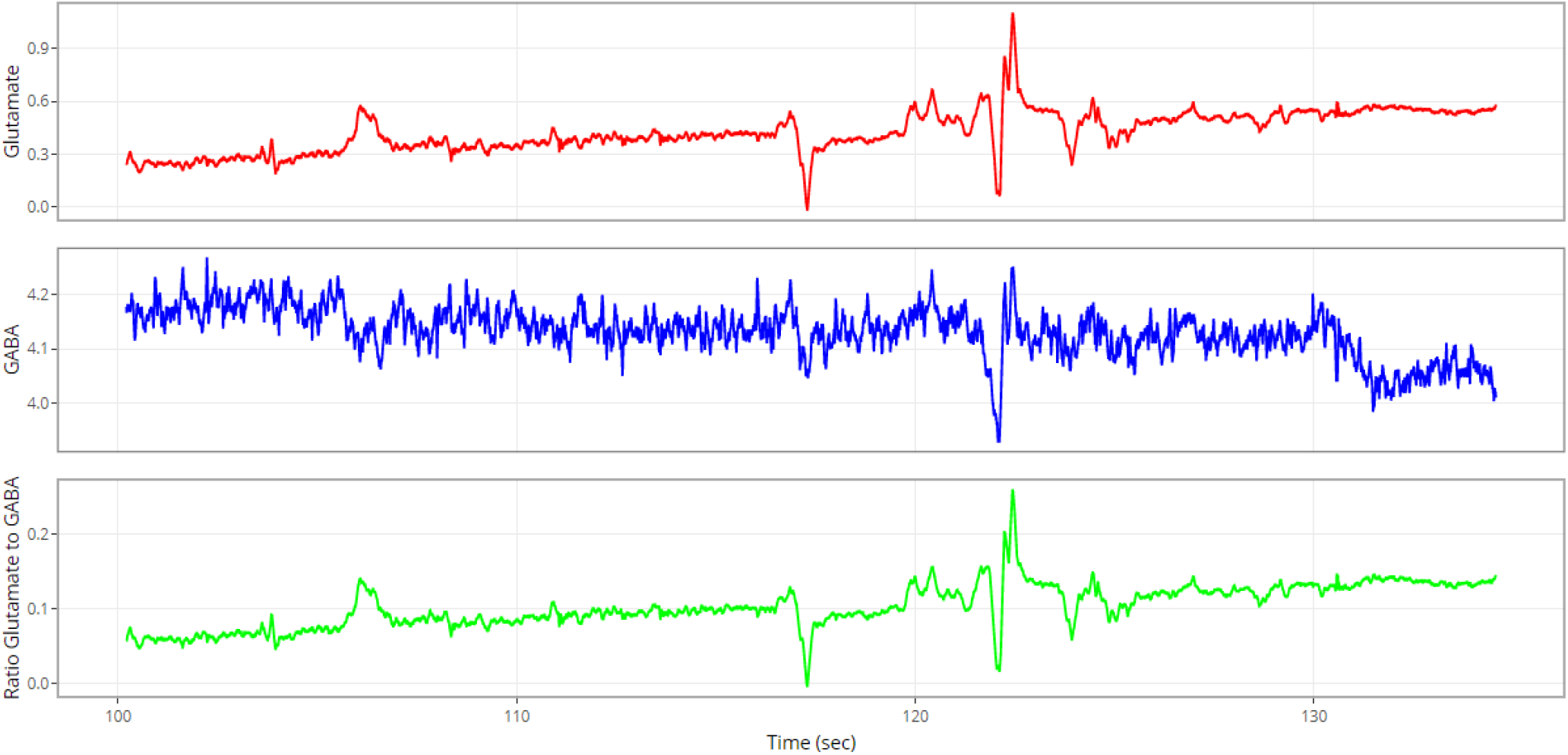
No subsampling and moving average by 10.

**Figure 14:**
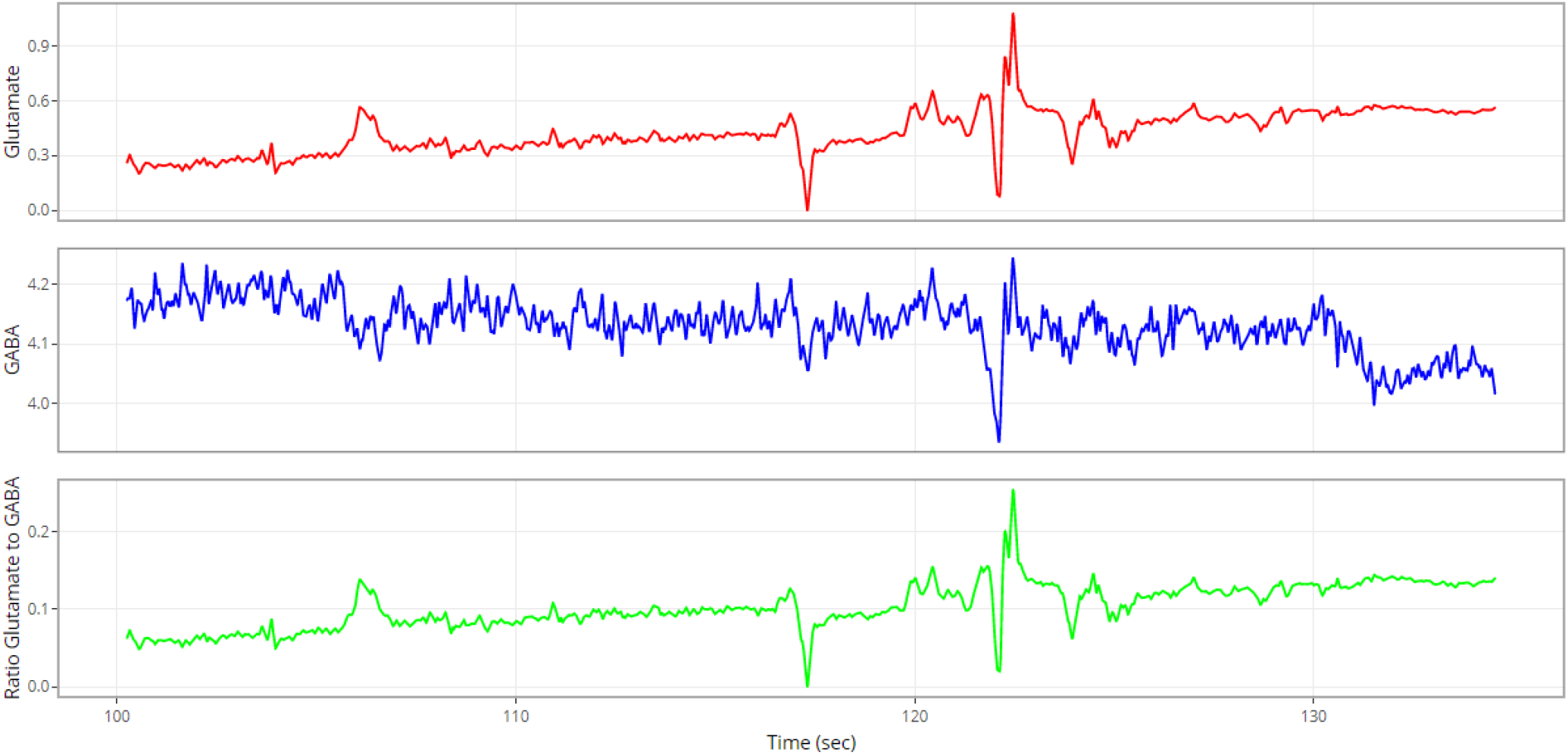
No subsampling and moving average by 50.

**Figure 15:**
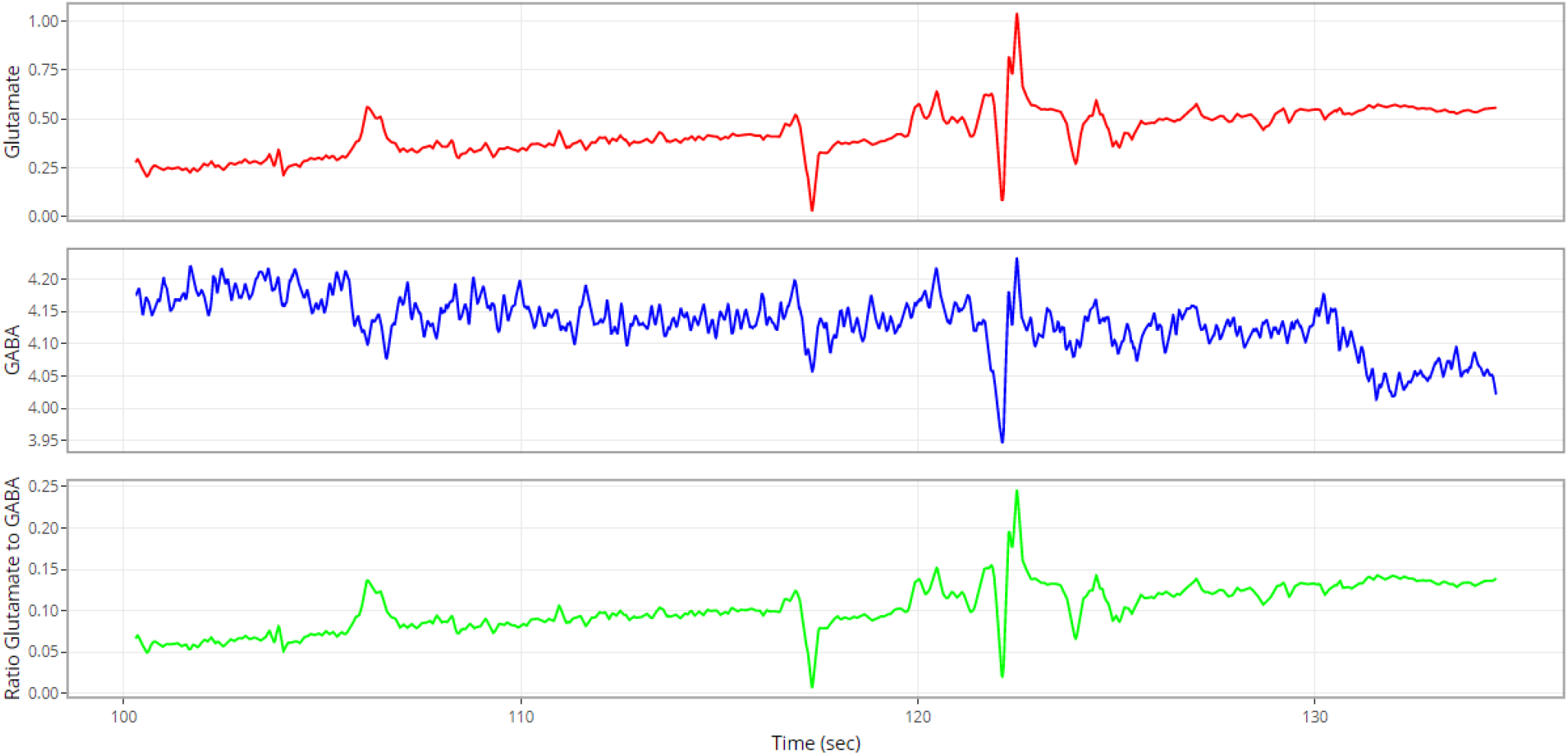
No subsampling and moving average by 100.

**Figure 16:**
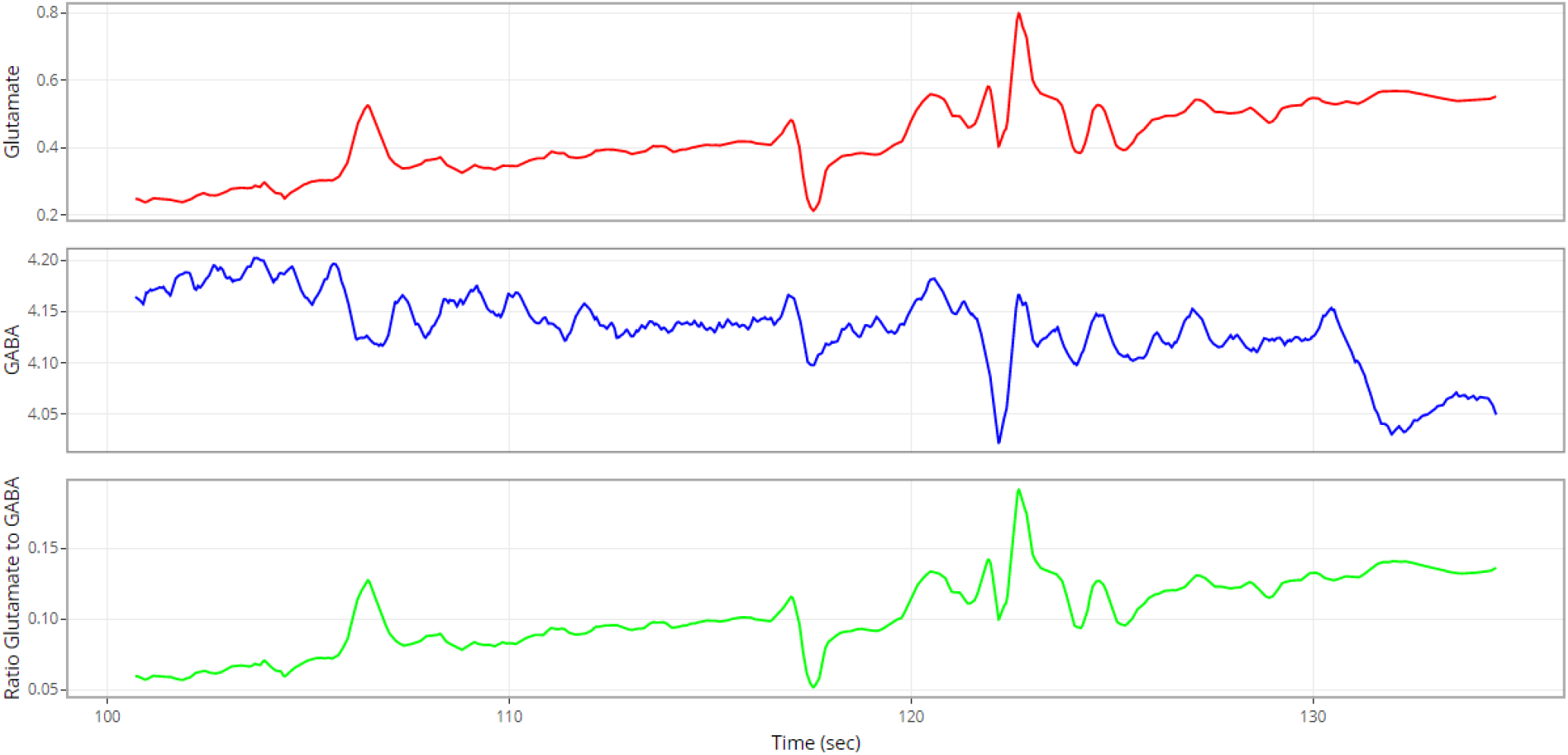
No subsampling; moving average by 500.

## 6 Conclusion

This report described a user-friendly, interactive visualization (implemented in R Shiny framework) to facilitate analysis and a better understanding of neurotransmitter data collected within the context of epileptic seizures.

Given the very high granularity of collected data (at millisecond level), it is challenging to use static visuals and/or tables for deeper data insights and features. Such challenges were greatly alleviated through an interactive visualizer (dashboard) which has ability to zoom out (for “big picture” analysis) and to zoom in (for a much more focused and targeted targeted analysis).

The visualizer is available at link https://kittyviz.shinyapps.io/GluGabaViz

Progress in development of real-time biosensors indicates the need for better analysis tools. The interactive visualizer described in the next section is particularly useful for analyzing what happens at higher frequencies in recordings.

## Notes

https://kittyviz.shinyapps.io/GluGabaViz

### Competing Interest Statement

The authors have declared no competing interest.

